# Early reduction in PD-L1 expression predicts faster treatment response in human cutaneous leishmaniasis

**DOI:** 10.1101/2020.02.21.959528

**Authors:** Nidhi S. Dey, Sujai Senarathna, Vijani Somaratne, Nayani Madarasinghe, Bimalka Seneviratne, Sarah Forrester, Marcela Montes De Oca, Luiza Campos Reis, Srija Moulik, Pegine Walrad, Mitali Chatterjee, Hiro Goto, Renu Wickremasinghe, Dimitris Lagos, Paul M. Kaye, Shalindra Ranasinghe

## Abstract

Cutaneous leishmaniasis (CL) is a chronic skin disease caused by *Leishmania* parasites and in Sri Lanka, CL is caused by *L. donovani.* Pentavalent antimonials (e.g. sodium stibogluconate; SSG) are first line drugs for CL, despite protracted and painful treatment regimens. Data from animal models indicate that the effectiveness of SSG requires drug-immune synergy, but mechanistic insight from patients is lacking. We studied whole blood and lesion transcriptomes from CL patients in Sri Lanka at presentation and during SSG treatment. In lesions, we identified differential expression of immune-related genes, including immune checkpoint molecules, after the onset of treatment whereas no differentially expressed genes were identified in whole blood. We confirmed reduced lesional PD-L1 and IDO1 protein expression on treatment in a second validation cohort, using digital spatial profiling and quantitative immunohistochemistry. Dual IHC-FISH revealed significantly higher expression of these immune checkpoint molecules on parasite-infected compared to non-infected lesional CD68^+^ monocytes / macrophages. Crucially, early reduction in PD-L1 but not IDO1 expression was predictive of rate of clinical cure and occurred in parallel with a reduction in parasite load. A multivariate cox proportional hazard model showed that patients with lower PD-L1 expression on treatment were more likely to cure earlier (HR= 4.88). Our data support a model whereby the initial anti-leishmanial activity of antimonial drugs alleviates checkpoint inhibition of T cell immunity, facilitating immune-drug synergism and clinical cure. Our findings demonstrate that PD-L1 expression can be used as an early predictor of clinical response to SSG treatment and support the use of PD-L1 inhibition as adjunct host directed therapy in Sri Lankan CL.

## Introduction

Up to one billion people are thought to be at risk of leishmaniasis, a group of diseases caused by infection with protozoan parasites of the genus *Leishmania* and transmitted by phlebotomine sand flies (1–3). Approximately 600,000 – 1 million new cases of CL occur, with a broad global distribution, often leading to stigma and reduced life chances and placing a burden on health services (4, 5). Treatment options for CL have changed little in over 70 years, since pentavalent antimonial drugs were first introduced, and there are scant new treatments on the horizon (6, 7). Sri Lanka is endemic for CL (8) with the first autochthonous case being reported in 1992 (9). Sri Lankan CL is caused by *Leishmania donovani* zymodeme MON-37(10–12), usually associated with visceral leishmaniasis in other endemic countries. Current treatment for CL in Sri Lanka involves weekly intra-lesional or daily intra-muscular administration of SSG, with or without cryotherapy, based on the site and size of the lesion and response to treatment. Cure often takes many months, and some patients may fail to respond completely or withdraw from treatment (13).

Most of our understanding of the host immune response in localised CL stems from experimental models of the disease, and human disease is much less well understood (14). Immune checkpoint molecules have been implicated in disease progression in pre-clinical models (15–23), but their role in human CL has not been explored. It is widely proposed that immune-drug synergy is required for fully effective treatment in leishmaniasis and that host directed therapy (HDT) may have an important future role in patient management (24–26), but few validated targets have emerged. Here, we searched for early correlates of treatment response that might be used to stratify patient response and shorten treatment duration. Our results indicate an intimate relationship between intracellular parasitism and immune checkpoint molecule expression, identify PD-L1 as a promising target for HDT in Sri Lanka.

## Results and Discussion

We reasoned that examination of the intra-lesional transcriptomic response early after the onset of therapy might reveal potential mechanisms underpinning immune-drug synergy and potential targets for HDT. We therefore conducted a targeted transcriptomic analysis of the lesion site in a test cohort of 6 patients with typical homogeneous nodulo-ulcerative CL lesions (3 females, 3 males; mean age ± standard deviation, 34 ± 11 years; (**Supplemental Figures 1-3** and **Supplemental Table 1**). Principal component analyses of lesion transcriptomic data showed separation for most patients comparing pre- to on-treatment samples (**Figure 1A**). Differential expression analyses comparing pre- and on-treatment transcriptomic profiles identified 120 differentially expressed genes (DEGs; FDR adjusted p-value<0.01; **Figure 1B**). To determine whether similar changes were also reflected in patient blood, we also conducted whole blood RNA-seq. No DEGs were identified (**Supplemental Figure 4**) suggesting that unlike CL caused by *L. braziliensis* (27), CL due to *L. donovani* in Sri Lanka is not accompanied by an overt systemic immune response.

**Figure 1.**
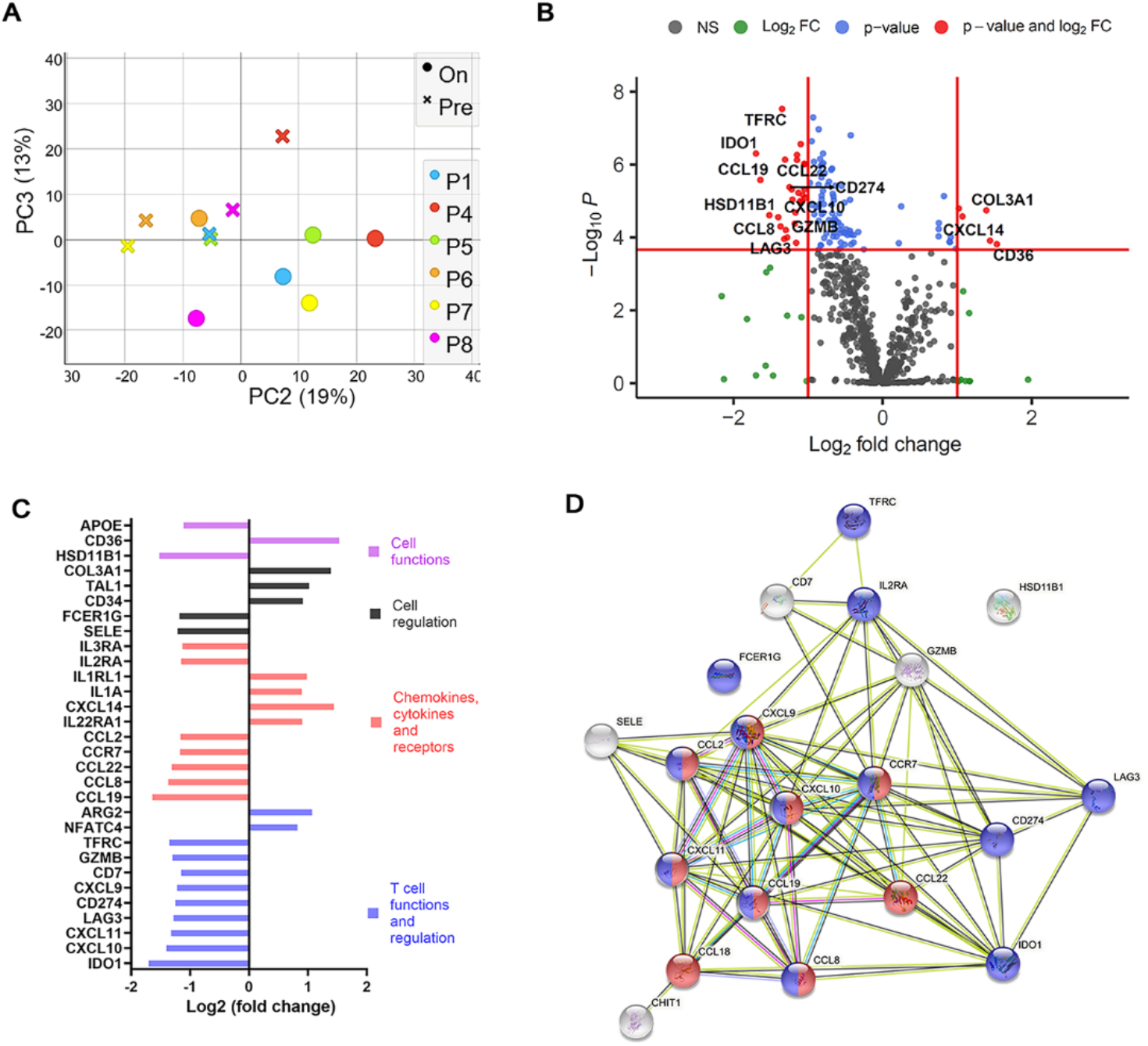
Differential expression and network analysis of genes regulated by drug treatment in lesions of Sri Lankan CL patients. Immune-targeted tissue transcriptomics was conducted on tissue sections from test cohort patients comparing transcriptomes at presentation and on treatment. (**A**) Principal component analysis was performed to show differences between pre- and on-treatment transcriptome of each patient based on 770 gene nCounter PanCancer Immunology Panel **(B)** Differentially expressed genes comparing pre-treatment biopsies with biopsies taken after two weeks on treatment with SSG. Cut off (red line) drawn at equivalent of adjusted p-value =0.01 and Log (Fold change) of 1**(C) T**op 30 genes that changed in expression on SSG treatment. **(D)** STRING protein-protein interaction network(28)(https://string-db.org) analysis of genes listed in **Supplemental Table 3** down-regulated on SSG treatment. Pathways represent GO:0072676, Lymphocyte migration (red spheres) and GO:0002684, positive regulation of immune system process (blue spheres). Top 20 genes are shown (Log2fold change ≥1.15) for clarity.

The majority of DEGs in lesion biopsies were downregulated (87%; 105/120) on treatment suggesting a reduction in inflammation with treatment (105 downregulated, 15 upregulated; **Figure 1B** and **Supplemental Table 3**. Genes for cellular functions and regulation, chemokines, membrane receptors, T cell function and regulation were amongst the top 20 DEGs **(Figure 1C**). Further, STRING analysis (28) identified Lymphocyte migration (GO: 0002687, FDR= 1.06E-14; including interferon inducible chemokines like *CXCL9, CXCL10, CXCL11*, *CCL19, CCL8)*) and regulators of immune response (GO: 0002684, FDR=1.94E-11; including *IDO1, LAG3* and *CD274/PD-L1*) as highly enriched pathways (**Figure 1D**). Transcripts of inflammatory mediators including *CXCL10*, *GZMB*, *CCL2* and *CCR7* (receptor for CCL19), previously shown to be associated with other forms of murine (29–31) or human CL (32–34) were also found to be downregulated with initiation of treatment (**Supplemental Table 3**).

We next conducted multiplexed antibody digital spatial profiling (35) for 59 immune targets, selecting regions of interest (ROIs) based on CD3^+^ and/or CD68^+^ expression (**Supplemental Figure 5** and **Figure 2A–F**). t-SNE dimensional reduction on a total of 33 ROIs analysed from three patients (P4, P6 and P7) (**Figure 2G**) indicated a degree of inter-patient heterogeneity in pre-treatment lesion protein profile, but with clear discrimination for each patient between pre- and on-treatment ROIs. IDO1 and PD-L1 as well as PD-1 were selectively reduced in expression upon treatment (**Figure 2H and I**). STRING analysis of all discoveries based on FDR (5%) also indicated significant enrichment in GO:002684, as well as a pathway associated with regulation of T cell activation (GO:0050863; **Supplemental Figure 6 A-B**).

**Figure 2.**
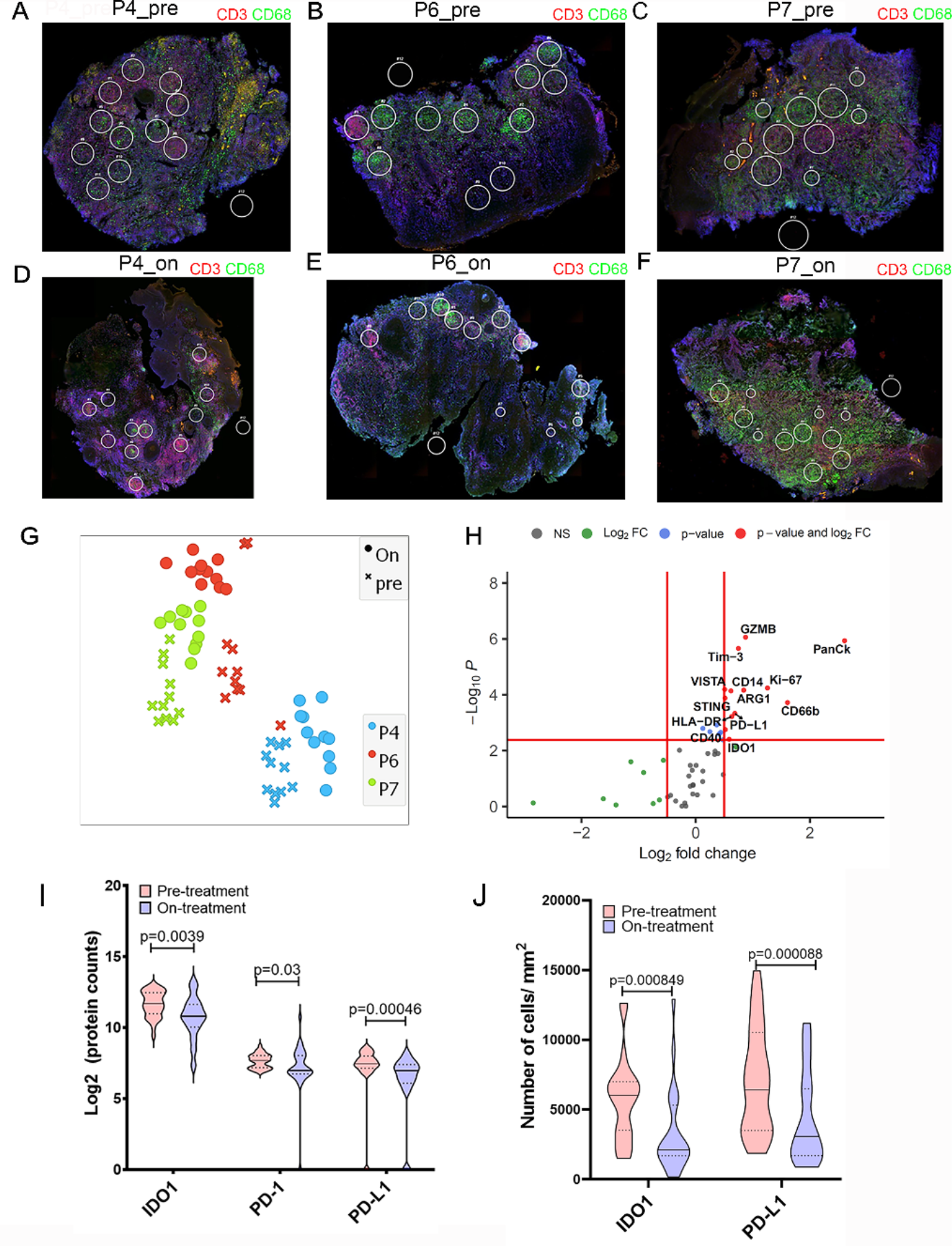
Digital Spatial Profiling of CL lesions. DSP was performed on tissue sections from test cohort individuals comparing ROIs from pre and on-treatment biopsies**. (A-F)** ROIs on CD3 and/or CD68^+^ rich areas from pre and on-treatment biopsies from patients P4, P6 and P7 (CD68, green; CD3, red; Syto13, blue). (**G)** t-SNE plot based on 20 PCA loadings coloured on patient ID. (**H**) Differential protein expression analysis comparing pre-treatment to on-treatment ROIs Red lines indicate adjusted p value cut off of 1% (Mann-Whitney test with FDR correction based on Benjamini, Krieger, and Yekutieli two stage set-up method) and and Log2FC = 0.5 (**I)** IDO1, PD-1 and PD-L1 expression in pre- and on-treatment ROIs. Mann Whitney rank test (n=33 ROIs). (**J**) IHC was performed on sections from patients pre and on-treatment from the validation cohort and quantitated using StrataQuest (see Methods) (n=23). Wilcoxon matched-pairs signed rank test.

As IDO1 and PD-L1 have been targeted in cancer immunotherapy and hold promise for drug re-purposing, we next sought to validate these findings by evaluating IDO1 and PD-L1 expression by quantitative IHC in an independent validation cohort of CL patients (5 females, 18 males; mean age ± standard deviation, 44 ± 11 years; time to diagnosis 7.76 ± 8.2 months; **Supplemental Figures 7 and 8** and **Supplemental Table 4**) sampled at baseline and after 4 weeks of treatment. Using an accepted cut-off of >5% of cells being positive (36), all patients (n=23) expressed IDO1 (Histochemical (H)-score (37) median = 81.2; range 16-165) and 20/23 patients had a reduction in the abundance of IDO1^+^ cells on treatment (H-score median = 32; range 1 – 171; p=0.0023; **Figure 2J**). All patients were PD-L1 positive at presentation (n=23; H-score median = 82.8; range 12-164) and 20/23 patients exhibited a reduction in the number of PD-L1 expressing cells on treatment (**Figure 2J**; H-score median = 36.7; range 12.3-36.7; p=0.0008). Collectively, these data indicate that IDO1 and PD-L1 are highly expressed in the lesions of Sri Lankan CL patients and that reduction in expression of these two checkpoint pathways represents an early response to SSG.

Though *in vitro* studies have indicated that intracellular parasitism by *Leishmania* could affect the expression of immune checkpoint molecules (38–40), this has not been established in situ during human disease. To address this question, we combined IHC with RNA-FISH (41) to identify *Amastin* transcripts (as a surrogate for viable amastigotes) with a bespoke StrataQuest image analysis pipeline (**Supplemental Figure 9, A-F**). In 7 patients studied that were *Amastin*^+^ at presentation (**Supplemental Methods, Supplemental Table 5**), PD-L1 expression co-localised with CD68^+^ macrophages (**Figure 3A, Supplemental Figure 10C**) and parasitized cells were both PD-L1^+^ and PD-L1^−^ (**Figure 3A**). We binned the Amastin^+^ PDL1^+^ and Amastin ^−^ PDL1^+^ cells based on PD-L1 mean fluorescent intensity (**Figure 3, B–D**) and found that cells containing abundant *Amastin* transcripts expressed more PD-L1 than cells with less or no *Amastin* transcripts **(Figure 3, B–E, Supplemental Figure 9, G-L** and **Supplemental Figure 10**). To independently corroborate this observation, we showed that the Sri Lankan strain of *L. donovani* was also capable of inducing up-regulation of PD-L1 expression on human monocyte-derived macrophages in vitro (**Supplemental Figure 11A-F**), as previously described for *L. major* (40). Similarly, IDO1 extensively co-localised with CD68^+^ cells (**Supplemental Figure 11A** and both IDO1^+^CD68^+^ and IDO1^−^CD68^+^ cells were infected (**Supplemental Figure 11B**). Using a similar gating strategy (**Supplemental Figure 11C-H**; n=3 patients), we found that cells with abundant *Amastin* transcripts expressed more IDO1 than those with fewer or no *Amastin* transcripts (**Supplemental Figure 11, I-K).** These data show that, although a notable population of uninfected CD68^+^ cells contribute to PD-L1 and IDO-1 expression within the CL lesion, intracellular parasitism leads to heightened expression of these checkpoint molecules by lesional monocytes and macrophages.

**Figure 3.**
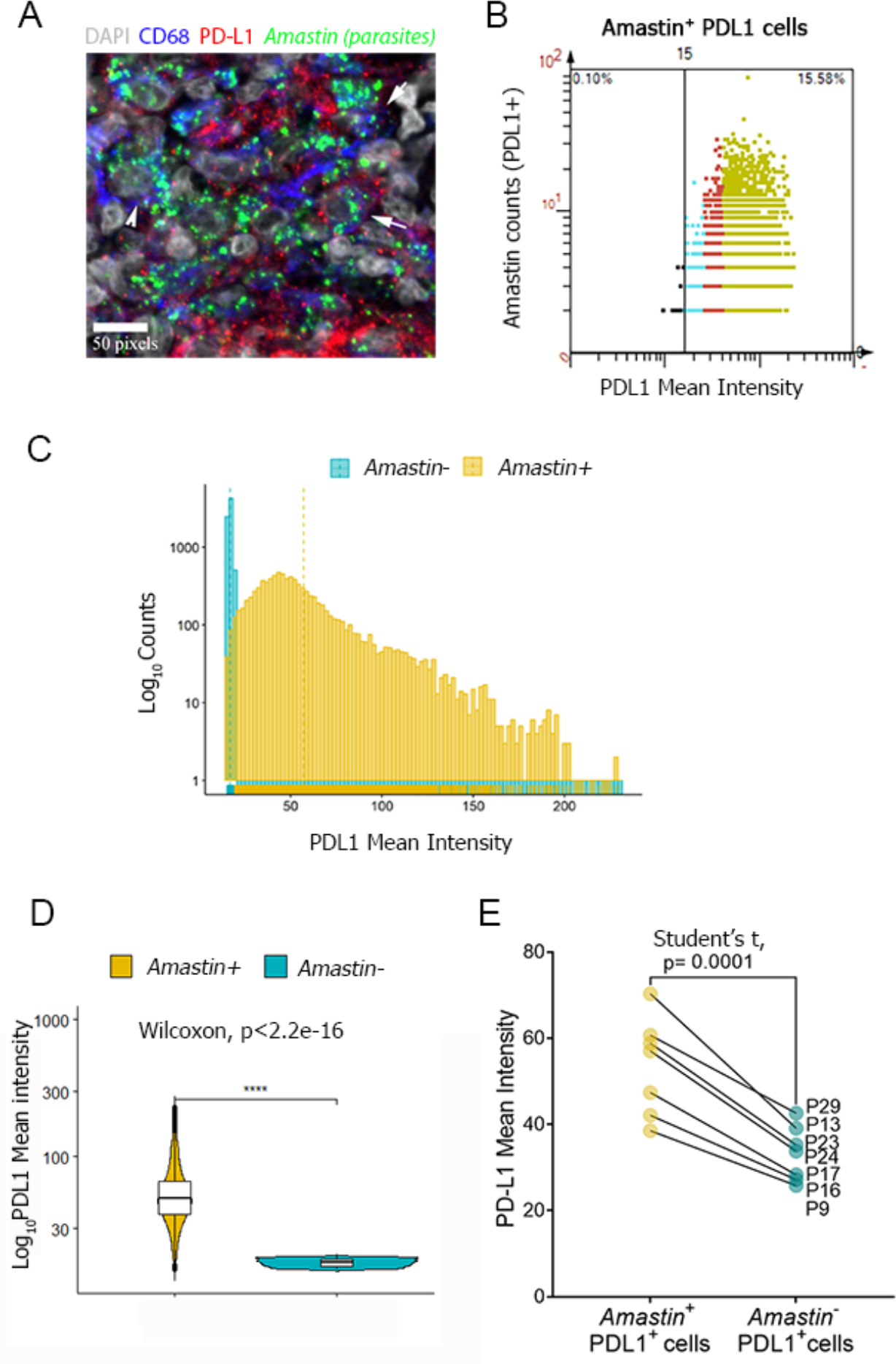
Imunofluorescence analyses of PD-L1 in infected and uninfected cells. Dual IHC-FISH using an *Amastin* probe was performed on pre-treatment sections of patients enrolled in the validation cohort. **(A)** Confocal image showing infection of PD-L1^+^CD68^+^ (arrows) and PD-L1^−^CD68^+^ (arrowhead) cells. Scale bar, 50 pixels **(B)** Relationship between PD-L1 expression and parasite burden (*Amastin* dot count). Scattergram from a representative patient (P24 at presentation) showing *Amastin*^+^ low (cyan), medium (red) and high (green) PD-L1 expressing cells with respect to parasite abundance. **(C)** Fluorescence intensity distributions of infected and uninfected PD-L1 cells. **(D)** Mean fluorescent intensity of PD-L1 expression on *Amastin*^−^ cells compared to *Amastin*^+^ cells from a representative patient P24. Dotted lines show median, upper and lower quantile. N=9159 parasite positive cells and N=41520 for parasite negative cells. Significance score was generated using Wilcoxon signed rank test. **(E)** PD-L1 expression on *Amastin*^+^PD-L1^+^ cells vs. *Amastin^−^*PD-L1^+^ cells (n=7 patients). Significance score was generated using Students two tailed paired t-test.

Finally, we tested whether reduction in IDO1 or PD-L1 expression early on during therapy could be used as a prognostic marker for treatment response. Patients with the greatest reduction in PD-L1 expression (i.e. greater than the geomean of the pre- : on-treatment expression ratio; n=12 patients) (**Figure 4A–B**) cured earlier than those that had lower or no reduction in PD-L1 expression (p=0.015). Patients with lower PD-L1 expression after 4 weeks of treatment (i.e. lower than the geomean of on-treatment expression; n=12 patients) also cured faster (p=0.0045; **Figure 4B**). We assessed the association of PD-L1 with disease cure rate using univariate Cox Proportional Hazard regression (**Supplemental Figure S13A**; Hazard Ratio (HR) = 3.96, p=0.008). Upon adjustment for age and gender of the participants, HR increased to 4.88 (p= 0.007; **Figure 4D**), indicating that patients that reduce PD-L1 expression most upon treatment are about 5 times more likely to cure earlier. Conversely, patients remaining parasite PCR^+^ at 4 weeks post treatment had significantly longer cure time (**Figure 4E**) and higher PD-L1 expression (**Figure 4F**). Surprisingly, even though intracellular parasitism was also associated with increased expression of IDO1, reduction in IDO1 expression, calculated as either pre:on-treatment expression ratio or IDO1 expression at 4 weeks (n=12 vs 11), did not correlate with cure rate (**Supplemental Figure 13 B and C**). Thus, the relationship between declining PD-L1 expression and rate of cure (**Figure 4, E–F**) appears selective with regards to other lesion-expressed checkpoint molecules.

**Figure 4.**
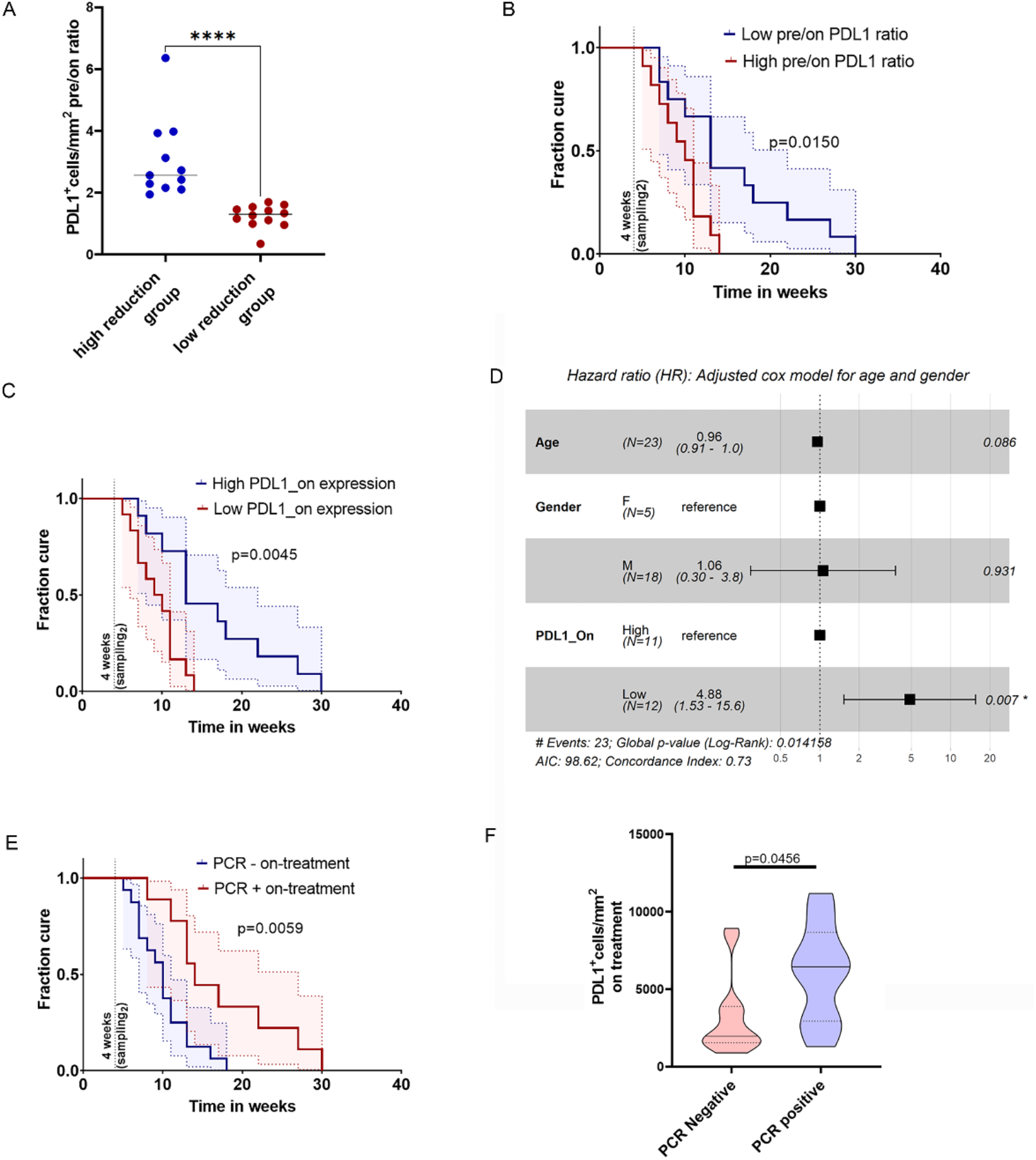
Clinical correlates of PDL1 reduction on treatment in CL patients. **(A)** Patients (validation cohort; n=23) were stratified based on high (>geomean value; n=11) and low (< geomean value; n=12) pre-: on-treatment expression ratio. **(B)** Kaplan-Meier curve based on pre-:on-treatment ratio of PD-L1 expression (high vs low). **(C)** Patients stratified based on on-treatment expression of PD-L1 (> geomean value; n=11 vs < geomean value; n=12). **(D)** Multivariate Cox Proportional Hazards model plotted as a forest plot. p-values for each covariate represent Wald statistic value and overall statistical significance is also indicated. Patients stratified by *LITS1* PCR status (n= 9 PCR^+^ vs n=14 PCR^−^ or ^+/−^ (equivocal)) on treatment. **(E)** PD-L1 expression in *LITS* PCR^+^ vs. PCR^−^ individuals on treatment. **(F-G)** Patients stratified for IDO1 based on pre-: on-treatment ratio (**F**) or on-treatment expression (**G**). Curves were compared using Log-rank (Mantel-Cox) test. Vertical line drawn in **B, C, E** on the X axis shows time when on-treatment biopsies were collected.

We conclude that expression of IDO-1 and PD-L1 immune checkpoint molecules is a common feature of Sri Lankan CL and that intracellular parasitism is associated with heightened expression of these immunoregulatory proteins in lesional macrophages. Tissue expression of both IDO1 and PD-L1 reduces significantly within a 2-4 weeks of treatment onset and well in advance of clinical cure, and a reduction in PD-L1 is associated with more rapid therapeutic response. The elevated expression of negative immune regulators on macrophages at the lesion site, as shown here, has clear parallels with tumour-associated macrophages (42) and extends our understanding of how *Leishmania* parasites influence the function of their host cell during human disease (43). Though longitudinal sampling of the same macrophage population is not possible, it seems likely that reduction of PD-L1 expression is facilitated by the leishmanicidal action of SSG, suggesting a model for drug-immune synergy whereby early rounds of SSG treatment reduce intracellular parasite burden leading to reduced checkpoint inhibition and re-engagement of T cell effector function. Our data, together with strong pre-clinical evidence of an inhibitory role of PD-L1 in various forms of leishmaniasis (17, 44, 45) supports the candidacy of PD-L1 blockade as an adjunct HDT in Sri Lankan CL. In addition, our data suggest the possibility that changes in PD-L1 expression early after treatment could also be used as a biomarker to trigger drug tapering or drug cessation.

## Methods

Information is provided in Supplemental methods.

### Study approval

The study was conducted in accords with the principles of the Declaration of Helsinki and was approved by the Ethical Review Committee of the Faculty of Medical Sciences, University of Jayewardenepura (Ref: 780/13 & 52/17) and the Department of Biology, University of York. Written informed consent was received from participants prior to inclusion in this study.

## Data availability

Digital Whole Slide Image files will be made available at https://leishpathnet.org. The raw data and read counts for RNA-seq data are available at the Gene Expression Omnibus under the accession X (to be provided on publication).

## Author Contributions

NSD design, experimental data analysis writing ms SS design, experimental, data analysis
VS experimental
NM experimental
BS experimental
SF data analysis
MMDO experimental
LR experimental
SM experimental
PW, conceptualisation design, funding
MC, HG, conceptualisation, design, funding
RW conceptualisation, design of study
DL conceptualisation design, data analysis, funding
PMK conceptualisation design, data analysis, funding, supervision
SR conceptualisation design, experimental, data analysis, funding, supervision

## Acknowledgements

The authors thank histopathologist, Dr. Pushpa Ilanngasinghe (Teaching Hospital Anuradhapura, Sri Lanka), phlebotomist, Dr. Dawei Chen (University of York), flow cytometry expert, Karen Hogg as well as other staff at Imaging and Cytometry Laboratory at BioSciences Technology Facility (University of York) for assistance with Strataquest and confocal analysis, technical support at Nanostring Technologies for DSP data acquisition, TissueGnostics for assistance with Strataquest, Centre for Genomic Research, University of Liverpool for processing samples for Nanostring transcriptomics and all members of the laboratory for their useful comments and suggestions. This work was supported by funding from the UK Medical Research Council / UK Aid Global Challenges Research Fund (MR/P024661/1 to PMK, SR, HG and MC) and Wellcome Trust Senior Investigator Award (WT104726 to PMK). The funders had no role in the design or conduct of the study of the decision to publish.

