## Supplemental methods and figures for "Early reduction in PD-L1 expression predicts faster treatment response in human cutaneous leishmaniasis"

**This file contains:**

**1. Methods**

**2. Supplementary Figures 1-13**

### Methods

#### Study settings

Test cohort sample collection was carried out at the Dermatology Unit, Teaching Hospital, Anuradhapura (THA) and Validation cohort sample collection was carried out at the District General Hospital Embilipitiya, Sri Lanka

#### Study design

This was designed as an observational longitudinal prospective study performed in Sri Lanka on two patient cohorts (Test and Validation). Individuals who were willing, had given written informed consent and having clinically highly suspected CL lesions were enrolled (age between 18-65 years). Clinical histories were taken, examinations were performed, slit skin smears and punch biopsies were obtained on day 0. After sample collection was completed, intralesional SSG was given on day 0 as a usual practice at the Dermatology clinics in Sri Lanka on clinically highly suspected CL lesions. Weekly intralesional SSG was given according to the guidelines given by Sri Lanka College of Dermatologists (Sirimanna et al., 2013). In the test cohort, follow-up examination and sample collection were carried out after 2 weeks and after 2 doses of SSG. 15 patients were enrolled, and sample collection was carried out from 29.10.2014-24.12.2014. Out of them PBMC could not be collected in 3 patients due to technical errors and therefore had to be excluded from the study group. From the remaining 12 patients, four did not report on the two-week follow-up day. Therefore, they were considered as lost to follow up. Out of followed up 8 patients, 7 completed treatment with compete clinical cure and 1 patient could not be contacted to obtain continuous treatment details and to assess complete clinical cure (P1; lost to final follow up) (**Figure S1**). In the Validation cohort follow-up examination and sample collection was performed after 4 weeks and after 4 doses of SSG. 30 patients were enrolled, and sample collection was carried out from 31.05.2018-20.09.2018. Four patients were excluded as they were recently diagnosed with diabetes mellitus and one was lost to follow up. The remaining 25 patients were followed-up until 6 months to assess complete clinical cure (24 patients had complete clinical cure by 6 months and one patient cured completely after 6.5 months). Of these, 2 patient sections were excluded from the analysis because of high background seen in IDO1 and PD-L1 staining (**Figure S5**). Biopsies were formalin fixed and embedded in paraffin (FFPE) and shipped to University of York (UoY) and subjected to immunological studies described here.

*Risk of bias assessment* No blinding method was included in the original design for this observational study. However, data & sample collection was carried out in Sri Lanka and analysis was done in the UoY by separate scientists. Therefore, scientist evaluating the samples was blinded to the clinical outcome at the time of analysis as clinical data was not sent to UoY until analysis was completed.

*Exclusion criteria* Individuals younger than 18 years and older than 65 years, pregnant women, patients with other concurrent/past illnesses or debilitation (diabetes mellitus, connective tissue disorders, immunosuppression, hypothyroidism, HIV/AIDS, Hansen's disease, peripheral vascular disease or lymphedema or varicosities in the affected limb, inflammatory dermatoses affecting the suspected lesion), patients with previous history and treatment of CL, individuals with lesions with signs of secondary bacterial infection with purulent exudates, patients who had a visit to a leishmaniasis endemic country at any point of life, indigenous population, persons with lesions in areas where biopsy cannot be performed, cases where treatment modality differed from intra-lesional SSG to intamuscular SSG, patients on medications that might impede or improve wound healing and scarring.

All patients were physically examined by a dermatologist at every visit to exclude or treat infected CL lesions, iatrogenic infections following sample collection or drug therapy.

**Patients and patient derived samples**

***Test cohort:*** Patients had 1-3 lesions (ulcers: 7/8; nodular ulcerative: 1/8) lesion with a mean time of diagnosis of 6.75 ± 5.8 months (**Supplemental Table 1)** and received intra-lesional SSG weekly (0.5-2ml/cm$^2$). Patients were diagnosed on basis of SSS, histopathology and/or clinical evaluation. In case of poor correlation between slit skin smear and histopathology, judgement was based on clinical assessment as it has been previously shown to be >90% accurate for diagnosis (1). P4 was included as it fit the clinical criteria for CL despite having no discernible amastigotes present (3) (**Table S1**). 3/6 of patients also received adjunct cryotherapy at a later stage of treatment when they were not responding sufficiently to intralesional SSG. At presentation, the amastigote density grading, based on slit skin smears (SSS) ranged from 0-6 (median =3) and was 0-3 after two weekly doses of SSG (median =2.5; not significant; Wilcoxon matched pairs signed rank test; **Supplemental Table 1**). Based on amastigote detection in H&E stained biopsy sections, 5/6 patients (**Supplemental Figure 3**) showed improvement in amastigote grade (1) after two rounds of SSG. 14.4 ± 4.15 doses of SSG were required to reach clinical cure (defined as complete re-epithelialization and flattening of edges; **Supplemental Table 1**). Mean area of ulceration of these individuals was 50 mm$^2$ (± SD, 35.1) and was calculated as detailed in section 'Lesional area calculation'.

***Validation cohort:*** Lesions were dry nodular ulcerative (n=8), dry ulcers (n=6), dry ulcerated plaques (n=5), wet ulcers (n=2), papular satellite lesions (n=2), nodule with satellite lesions (n=1), ulcerated plaque with satellite lesions (n=1) with mean ulcerated area of 51.13 mm$^2$ (±

SD, 81.1) and mean area of induration of 1430.2 ( ± SD, 1966.9). In addition to clinical assessment, CL was diagnosed by slit skin smears (SSS) (18/23) and PCR (23/23). Punch biopsies were taken from these patients at baseline and after 4 weeks of treatment with weekly dose of 1 ml/cm$^2$ intralesional SSG. At presentation, the amastigote density grading, based on slit skin smears (SSS) ranged from 0-5 (median =2) and was 0-5 after two weekly doses of SSG (median =0; Wilcoxon matched pairs signed rank test; p= 0.0001; **Supplemental Table 5**).

***Slit skin smears (SSS):*** Tissue scrapings from a 3mm superficial nick from the active edge of the lesions were used to prepare smears on slides, stained with Giemsa and examined under oil immersion microscopy for the presence of amastigotes. Parasite density was graded from 0 to 6+ according to WHO guidelines for VL(1): 0—no parasites per 1000 high power fields (HPF: x 1000 magnification); 1+: 1–10 parasites per 1000 HPFs; 2+: 1–10 parasites per 100 HPFs; 3+: 1–10 parasites per 10 HPFs; 4+: 1–10 parasites per HPF; 5+: 10–100 parasites per HPF; and 6+: > 100 parasites per HPF.

***Punch Biopsy:*** A 3mm diameter full thickness punch biopsy was taken from the edge of the lesion under local anaesthesia, transported in formol saline and then fixed in paraffin blocks and used for H&E (Test cohort only) and FISH + IF & trancriptome analysis studies. Simultaneously in the Validation cohort an 2mm diameter punch biopsy was taken for PCR

***Lesional size calculation***

Lesion size was analysed as a measure of total area of the lesion (ulcerated and indurated). Ulcer diameters (D1, D2), induration diameters (D3, D4) and width of margin between ulcer and induration (W1) were measured and total area was calculated as below (2)

$$(D1+D3+W1)*(D2+D4+ W1)$$

**PCR**

From each lesion in the Validation cohort, another 2 mm diameter full thickness punch biopsy sample was taken from the active edge, stored in RNA later at -20˚C and was subjected to DNA extraction using QIAGEN DNeasy Blood & Tissue Kit. A volume of 2µl of this extracted DNA was amplified with 100 pmols of previously described(3) set of primers LITSR/L5.8S that amplifies a 320 bp fragment of ITS1 region of Leishmania genus-specific DNA in the presence of 1.5 mM $MgCl_2$, 200 µM of each deoxynucleotide triphosphate, 25 units/ml Taq DNA polymerase, reaction buffer (pH 9.0) (Promega) in a final volume of 10 µL. The PCR amplification cycles consisted of an initial denaturation at 95°C for 2 min, followed by 34 cycles of denaturation at 95°C for 20 s, annealing at 53°C for 30 s, and extension at 72°C for 1 min, with a final extension of 72°C for 6 min (3, 4). Extracted DNA from a pure culture was the positive control and no DNA sample was the negative control. A volume of 5 µL from the PCR product was run at 100 V for 45 min on a 1.75% (w/v) wide range agarose gel stained with 0.2 µg/mL ethidium bromide (EtBr) in 1X tris-acetate-EDTA buffer. The image was visualised under UV light and captured by a computerised gel documentation unit (Quantum ST5; Vilber Lourmat, Germany).
LITSR (forward): 5'-CTGGATCATTTTCCGATG-3' L 5.8S (reverse): 5'-TGATACCACTTATCGCACTT-3'

**RNA isolation**

The total RNA was extracted from formalin-fixed paraffin-embedded (FFPE) patient samples using RNeasy FFPE Kit (Qiagen) as per manufacturer's protocol. Briefly, 2 10µm sections were cut from each block and put into deparaffinization buffer (Qiagen). Sample lysis was done with Proteinase K digestion for 15 minutes. After lysis, samples were incubated at 80ºC for 15 minutes followed by 15 minutes of DNase treatment. Finally, concentrated RNA was purified using RNeasy MinElute spin columns, and eluted in a volume of 14–30 µl.

**NanoString nCounter assay**

RNA quality and size of RNA fragments for nanostring assay was assessed using the Agilent 2100 Bioanalyser at Technology Facility, University of York. One-hundred nanograms of RNA was used for the analysis using the nCounter PanCancer Immune Profiling Panel (XT-CSOMIP1-12, NanoString Technologies(5)). Processing of samples and data collection were done at Centre for Genomic Research, University of Liverpool. Data analysis was performed using the nSolver Advanced Analysis Software (NanoString Technologies) v2.0. Filtering of samples using quality control criteria was performed and data was corrected for background expression. 37 gene probes were excluded for falling below background. Gene counts were further normalised by scaling with the geometric mean of the built-in housekeeping control gene probes (total HK genes in the panel were 40) for each sample according to the manufacturer's recommendations. HK gene selection was based on geNorm pairwise variation statistic within the nsolver 4.0 data analysis suite. Of 40 HK genes, 32 were used for normalisation.

**Bioinformatics analysis of the NanoString nCounter assay**

Log2-transformed normalised data was used to plot volcano plot of differentially expressed genes in R using Enhanced volcano package. Volcano Plot filtering (fold change $\geq 1$, adjusted $P<0.01$) was used to identify differentially expressed genes with statistical significance between the two groups. PCA analysis was done in Orange data mining tool (v 3.29). Wilcoxon paired signed rank test was applied to compare normalized expression values between groups with FDR correction (1%) based on Benjamini, Krieger, and Yekutieli two stage set-up method. Physical and functional interactions between proteins were determined using the STRING platform at a medium confidence score of 0.4.

**RNAseq**

Adaptors were removed using cutadapt v 1.8.3 using the parameters –a , -A and –p to remove universal adaptors from both 5' and 3' ends of reads (6). Unpaired reads were removed, and reads were further trimmed and scored for quality using sickle v 1.33 (https://codeload.github.com/najoshi/sickle/tar.gz/v1.33). Reads were then aligned using STAR aligner v 2.5.1b using the sjdbOverhang 150 to the human reference version GRCH38.p5 and annotation version GRCh38.84 (7, 8). Reads were assigned to the annotations in the reference using the union algorithm in HTseq-count(9). Unexpressed genes and those with logCPM lower than 1 were removed. DE expressed genes were identified using the R package edgeR (10). edgeR package was also used to normalised the gene counts, and pairwise comparisons were performed using a generalised linear model between the post on and pre -treatment groups, and using a FDR cut off of 0.05. PCA analysis was performed on all features using Orange data mining tool v 3.29. Volcano plot was generated using enhanced volcano packge in R [5].

**Nanostring DSP**

4µm thick sections were used for DSP analysis of 59 immune parameters (**Figure 2**) through Nanostring Technology Access Program (TAP) on FFPE skin biopsies. Slides were stained with CD3 and CD68 as morphological markers and with 62 oligo-nucleotide conjugated antibodies on three patients P4, P6, P7 at presentation and on treatment. Amongst 12 ROIs per sample, 11 ROIs were created and distributed uniformly on CD68+ macrophage and CD3 rich T cell areas and one was created outside the tissue on glass. Data were normalised to positive ERCC controls and glass values (ROIs created on glass and not the tissue) were deducted. Data was background thresholded to geomean of isotype controls (any value >GM of isotype controls /ROI were floored to zero) and scaled to geomean of housekeeping controls (S6, GAPDH and Histone H3). Data was plotted from all 33 ROIs on pre- and on-treatment sections and Mann-Whitney test was used to calculate significance scores with FDR correction (1%) based on Benjamini, Krieger, and Yekutieli two stage set-up method. t-SNE plots were generated from 20 PCA loadings (98.5% variability) in Orange data mining tool (v 3.29) with two perplexity values (50 and 500) to reveal global relationship between data points. Data was normalised column wise by subtracting column mean and dividing by standard deviation. Volcano plot was generated in R-4.1.0 using Enhanced Volcano package. Fold change was calculated by dividing average of pre-treatment counts/ feature with average of on-treatment counts/feature. Log2 transformation was then formed. P-values were adjusted for multiple comparisons using FDR correction (1%) based on Benjamini, Krieger, and Yekutieli two stage set-up method.

**IF and FISH**

4 µm sections from FFPE blocks of skin biopsies were used for all analyses. Skin sections were stained with H&E (Sigma Aldrich) and were imaged using Zeiss AxioScan.Z1 slide scanner. For immunofluorescence and RNAScope FISH assay, paraffin was removed from the formalin-fixed sections using Histo-clear followed by washes in alcohol and water. Antigens were unmasked by mild boiling in RNAscope® Target Retrieval Reagent (15 minutes), peroxidase treatment followed by treatment with RNAscope® Protease Plus Reagent for 30 minutes at $40^{0}$C. Parasites were detected by FISH using a 13ZZ probe against *L.Infantum*-amastin targeting 2-635 of LinJ.34.1010 (having 85% sequence homology with

LmjF.34.0500) for 2 hours at $40^0$C and developed using manufacturer's protocol. For IF-FISH dual staining, FISH was done first, followed by IF.

For IF, following a few washes with TBST (0.05% TBS Tween20), blocking was done with 5% (v/v) normal donkey serum, 5% (v/v) normal goat serum and 5% BSA (w/v) for 1 hr at RT. Sections were then probed with, mouse anti-human CD3 (1:10, Abcam USA, ab17143), rabbit anti-IDO1 (1:80, Cell Signaling Tech., D5J4E); rabbit anti-PD-L1 (1:150, Cell Signaling Tech., E1L3N), rabbit anti-human CD68 (1:800, Abcam USA, ab213363), mouse anti-human CD68 (1:100, Abcam USA, ab955), rabbit IgG isotype control (concentration same as the primary, Abcam USA, ab172730, mouse IgG1 isotype control (BioLegend USA, 401401) overnight at $4^0$C. Primary antibodies were detected by goat anti-mouse IgG (H+L) cross-adsorbed secondary antibody, Alexa Fluor 488 (Thermo Fisher Scientific, USA, and A11001) and goat anti-rabbit IgG (H+L) highly cross-adsorbed secondary antibody, Alexa Fluor 555 (Thermo Fisher Scientific, USA, A21429). Sections were counterstained with DAPI and mounted in Fluoroshield™ histology mounting medium.

**Image acquisition and analysis.**

Images were acquired using Zeiss AxioScan.Z1 slide scanner, Zeiss LSM 880 with Airyscan on an Axio Observer.Z1 invert and Zeiss LSM 710 on an AxioImager. M2. Identical exposure times and threshold settings were used for each channel on all sections of similar experiments. Quantification of IDO1, PD-L1, CD3 *Amastin*, and CD68 was performed using StrataQuest Analysis Software (TissueGnostics). In brief, for IDO1, CD68 and PD-L1 cells, the software segmented nuclei on the basis of the signal from the DAPI channel, then built and expanded a mask over staining of IDO-1/ PD-L1. A cut off was applied on IDO-1/PDL-1 mean intensities based on visual inspection of the tissue to delineate IDO1/ PD-L1 $^{+/-}$ cells.

On the IDO1 generated mask, algorithm searched for colocalization of a similar mask for CD68 signal for IDO1 CD68 double positive cells as well as IDO1$^{+}$CD68$^{-}$ cells. A separate layer was created to extract the information of *Amastin* positive dots in all patients. *Amastin*$^{+}$ dots were detected in 3/6 (test cohort) and 7/25 (validation cohort) of patients (within the limits of detection in thin sections; 4μ). Number of *Amastin* dots in IDO1$^{+}$ and PD-L1$^{+}$ cells were computed and plotted in histograms (**Supplemental Figure 8f** and **Supplemental Figure 10f**). A cut-off of one was applied to number of *Amastin* dots to look at IDO1$^{+}$*Amastin*$^{+}$/ PD-L1$^{+}$*Amastin*$^{+}$ nuclei. More histograms were plotted for PD-L1 and IDO1 mean intensity but gated onto the right and left quadrant of histograms for *Amastin* dots to get information of parasite positive and negative cells, respectively (**Figure 3c-d, 4c-d, Supplemental Figure 8f g-l, Supplemental Figure 11**). Gates were then generated on histograms of IDO1 and PD-L1 mean intensities to look at low, medium and highly labelled cells and were coloured as cyan, red and orange/green, respectively in separate scattergrams for *Amastin*$^{+}$/*Amastin*$^{-}$ PD-L1 and *Amastin*$^{+}$/*Amastin*$^{-}$ IDO1 cells. Histogram in Figure 3C and violin plot in Figure 3E were generated in R using ggplot2 package. Histochemical score (H-score(11)) was used to assay expression levels of IDO1 and PD-L1. H-score was calculated based on the formula = 0 x (% of IDO1/PD-L1$^{-}$ ) + 1 X (% of weakly labelled IDO1/PD-L1$^{+}$) + 2 X (% of moderately labelled IDO1/PD-L1$^{+}$) + 3 X (% of strongly labelled cells). Positive expression of PD-L1 and IDO1 was defined by positive staining in >5% of cells(12).

**Infection of hMDM1 and hMDM2**

Promastigotes of SL-CL clinical isolate (*L.donovani* MON-37) were cultured in M199 (Sigma) with 10% (v/v) HiFCS,100 units of penicillin, 0.1 mg streptomycin per mL (PenStrep; Sigma), 1M HEPES, 10mM adenine, 0.25% Hemin and 0.25 mg/ml of biopterin. . Promastigotes were incubated at 26 °C; the maximum in vitro passage number used was 8.

PBMCs isolated from 30mL of peripheral blood from healthy UK volunteers on day 0 (n=3 for first experiment and n=5 for second experiment) using Ficoll centrifugation method. CD14+ cells were MACS purified (Miltenyi, 130-050-201) from PBMCs and differentiated into M1 or M2 with 10ng/mL of GM-CSF (Miltenyi, 130-093-862) and 30ng/mL M-CSF (Miltenyi, 130-093-963) supplemented complete media (CM; RPMI1640, 10% FCS, 2 mM l-glutamine, 50 µM β-mercaptoethanol, 100 U/mL penicillin, 100 µg/mL streptomycin, 1 mM HEPES) respectively and incubated at 37ºC + 5%CO2 for 5 days. On D5, human monocyte derived macrophages (hMDMs) were detached, counted and seeded in equal numbers in eppendorfs with MOI of 10 with either *L.donovani* zymodeme MON-37 (CL) or *L.major* Friedlin (L.mf) stationary-phase CFSE labelled promastigotes in for 24 hours. *L. major* (MHOM/IL/81/Friedlin strain) strain was used as a positive control for PDL1 expression in leishmania infected HMDMs as previously shown (13). For CFSE labelling of parasites, 5E+06 d7-9 old parasites were incubated with 10mM CFSE for 12 minutes in the dark at room temperature. Excess CFSE was washed with complete media twice. Parasites were counted and seeded with macrophages. On D6, after 24 hours of infection, extracellular parasites were then removed by centrifugation and washing with complete media and assayed for PDL1 expression through flow cytometry. For flow cytometry analysis, cells were labelled with fluorescent antibodies for half an hour at 37C. List of antibodies used in this study are listed below.

| Supplier | Item | Clone | Cat # |
|---|---|---|---|
| BD Biosciences® | PE Mouse Anti-Human CD14 | M5E2 | 557154 |
| BD Biosciences® | BV650 Mouse Anti-Human CD14 | M5E2 | 563420 |
| BD Biosciences® | BUV395 Mouse Anti-Human CD163 | GHI/61 | 745572 |
| BioLegend® | PE/Cyanine7 anti-human CD274 (B7-H1, PD-L1) | 29E.2A3 | 329718 |
| BioLegend® | APC/Cyanine7 anti-human HLA-DR | L243 | 307618 |
| BioLegend® | PE/Cyanine7 Mouse IgG2b, κ Isotype | MPC-11 | 400325 |
| Thermo Fisher | IR885/LIVE/DEAD™ Fixable Near-IR stain | - | L34975 |
| BioLegend® | BV405/ Zombie Aqua™ Fixable Viability Kit | - | 423101 |

**Multivariate Cox proportional hazard model**

Multivariate as well as univariate hazard models were generated in R using survival and survminer packages. Statistical scores correspond to Wald's statistic value. A univariate model was generated to assay effect on PDL1 on treatment values on cure curves. This was then adjusted for age and gender of participants and hazard ratio was calculated and plotted as a forest plot in Figure 4D

**Statistics**

Statistical analysis was performed in GraphPad Prism (version 9) and in R 4.1.0. A P value less than 0.05 (*) was considered significant.


1. WHO Expert Committee on the Control of the Leishmaniases and World Health Organization. Control of the leishmaniases: report of a meeting of the WHO Expert Commitee on the Control of Leishmaniases, Geneva, 22-26 March 2010. https://apps.who.int/iris/handle/10665/44412. Updated 2010.
2. Olliaro P, Vaillant M, Arana B, Grogl M, Modabber F, Magill A, et al. Methodology of clinical trials aimed at assessing interventions for cutaneous leishmaniasis. *PLoS Negl Trop Dis.* 2013;7(3):e2130.
3. el Tai NO, Osman OF, el Fari M, Presber W, and Schonian G. Genetic heterogeneity of ribosomal internal transcribed spacer in clinical samples of Leishmania donovani spotted on filter paper as revealed by single-strand conformation polymorphisms and sequencing. *Trans R Soc Trop Med Hyg.* 2000;94(5):575-9.
4. Ranasinghe S, Wickremasinghe R, Hulangamuwa S, Sirimanna G, Opathella N, Maingon RD, et al. Polymerase chain reaction detection of Leishmania DNA in skin biopsy samples in Sri Lanka where the causative agent of cutaneous leishmaniasis is Leishmania donovani. *Mem Inst Oswaldo Cruz.* 2015;110(8):1017-23.
5. Goytain A, and Ng T. NanoString nCounter Technology: High-Throughput RNA Validation. *Methods Mol Biol.* 2020;2079:125-39.
6. M. M. Cutadapt removes adapter sequences from high-throughput sequencing reads. *EMBnetjournal* 2011;17.
7. Dobin A, Davis CA, Schlesinger F, Drenkow J, Zaleski C, Jha S, et al. STAR: ultrafast universal RNA-seq aligner. *Bioinformatics.* 2013;29(1):15-21.
8. Yates AD, Achuthan P, Akanni W, Allen J, Allen J, Alvarez-Jarreta J, et al. Ensembl 2020. *Nucleic Acids Res.* 2020;48(D1):D682-D8.
9. Anders S, Pyl PT, and Huber W. HTSeq--a Python framework to work with high-throughput sequencing data. *Bioinformatics.* 2015;31(2):166-9.
10. Team RC. R: A language and environment for statistical computing. *R Foundation for Statistical Computing, Vienna, Austria.* 2020.

11. Igarashi T, Teramoto K, Ishida M, Hanaoka J, and Daigo Y. Scoring of PD-L1 expression intensity on pulmonary adenocarcinomas and the correlations with clinicopathological factors. *ESMO Open.* 2016;1(4):e000083.
12. Powles T, Eder JP, Fine GD, Braiteh FS, Loriot Y, Cruz C, et al. MPDL3280A (anti-PD-L1) treatment leads to clinical activity in metastatic bladder cancer. *Nature.* 2014;515(7528):558-62.
13. Filippis C, Arens K, Noubissi Nzeteu GA, Reichmann G, Waibler Z, Crauwels P, et al. Nivolumab Enhances In Vitro Effector Functions of PD-1(+) T-Lymphocytes and Leishmania-Infected Human Myeloid Cells in a Host Cell-Dependent Manner. *Front Immunol.* 2017;8:1880.

**Supplemental Figure 1. CONSORT flow diagram**

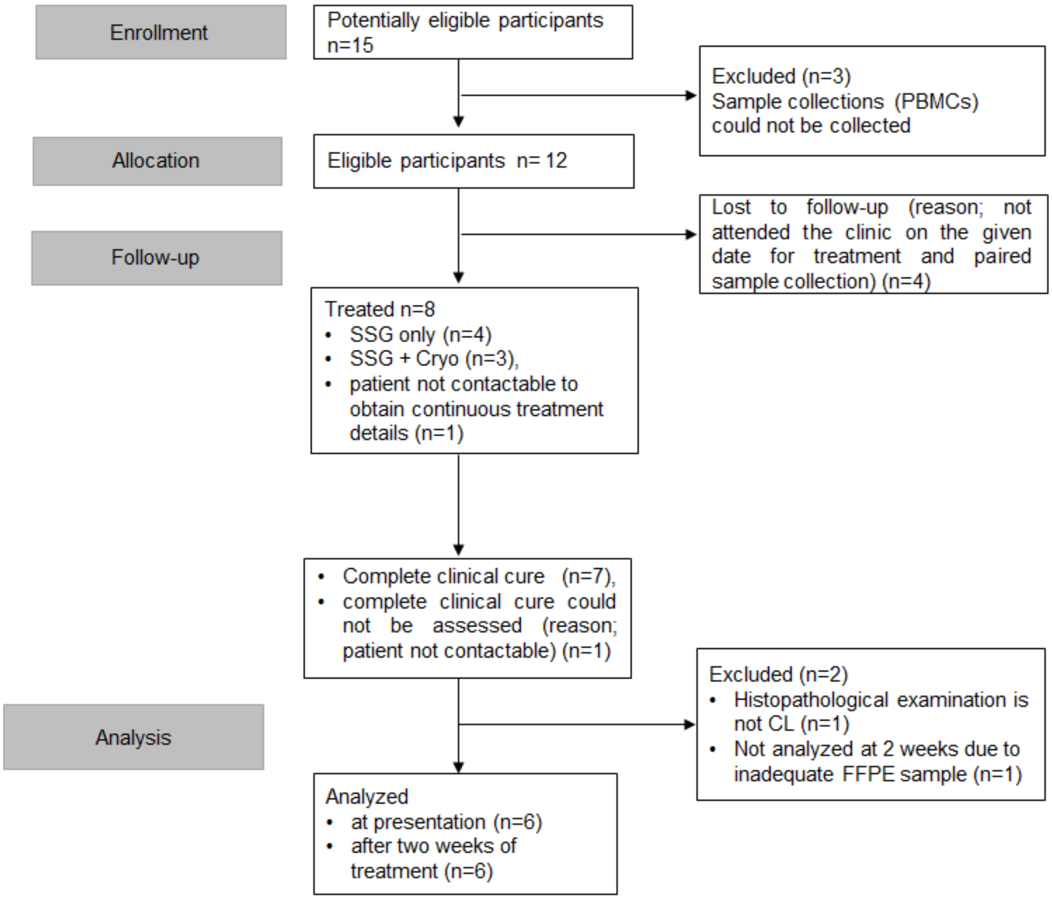


Test cohort patients that were screened, treated, followed up in early assessment, completed the study and included in the final analysis. For more details on the patients refer to Supplemental Table 1. PBMCs: Peripheral Blood Mononuclear Cells; SSG: Sodium Stibogluconate; Cryo: Liquid nitrogen cryotherapy.

**Supplemental Figure 2. CL lesions from test cohort patients**

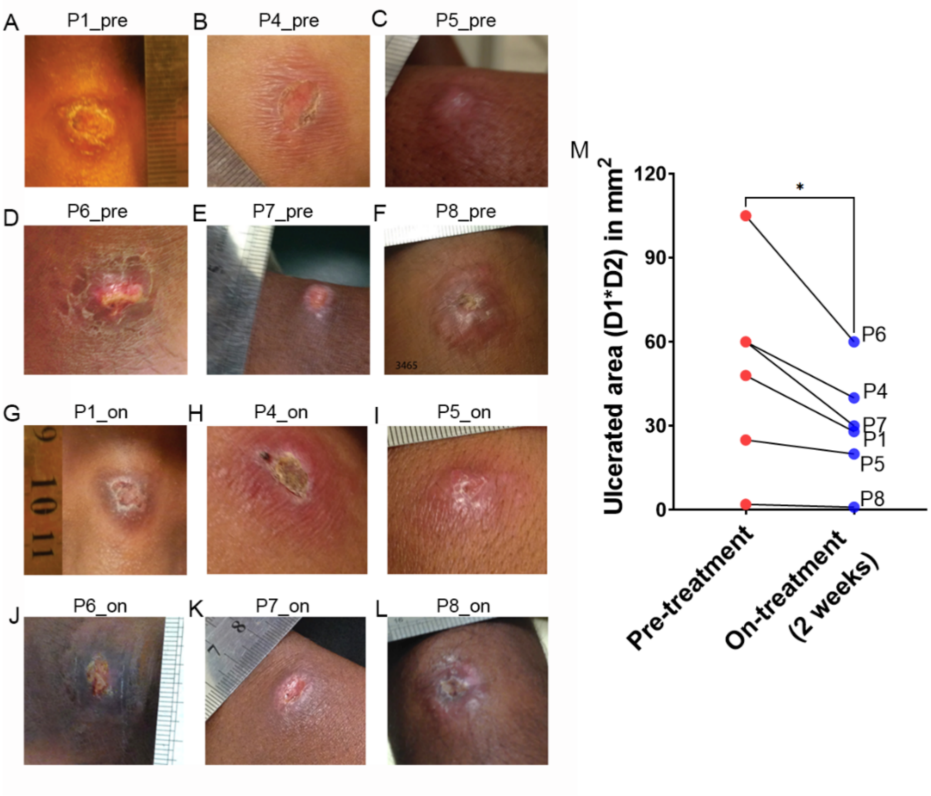


Lesions in patients were photographed at presentation and 3mm tissue biopsy were taken from the border of the lesion. **(A-L)** Lesion photographs from patients P1 (**A**), P4 (**B**), P5 (**C**), P6 (**D**), P7 (**E**) and P8 (**F**) at presentation and during treatment (**G-L)**. Comparison of total ulcerated area (Diameter 1*Diameter 2, when D1 and D2 were measured perpendicular to each other) per patients at presentation and after 2 weeks. Wilcoxon matched pairs signed rank test was used to calculate significance.

**Supplemental Figure 3. Histopathology of test cohort**

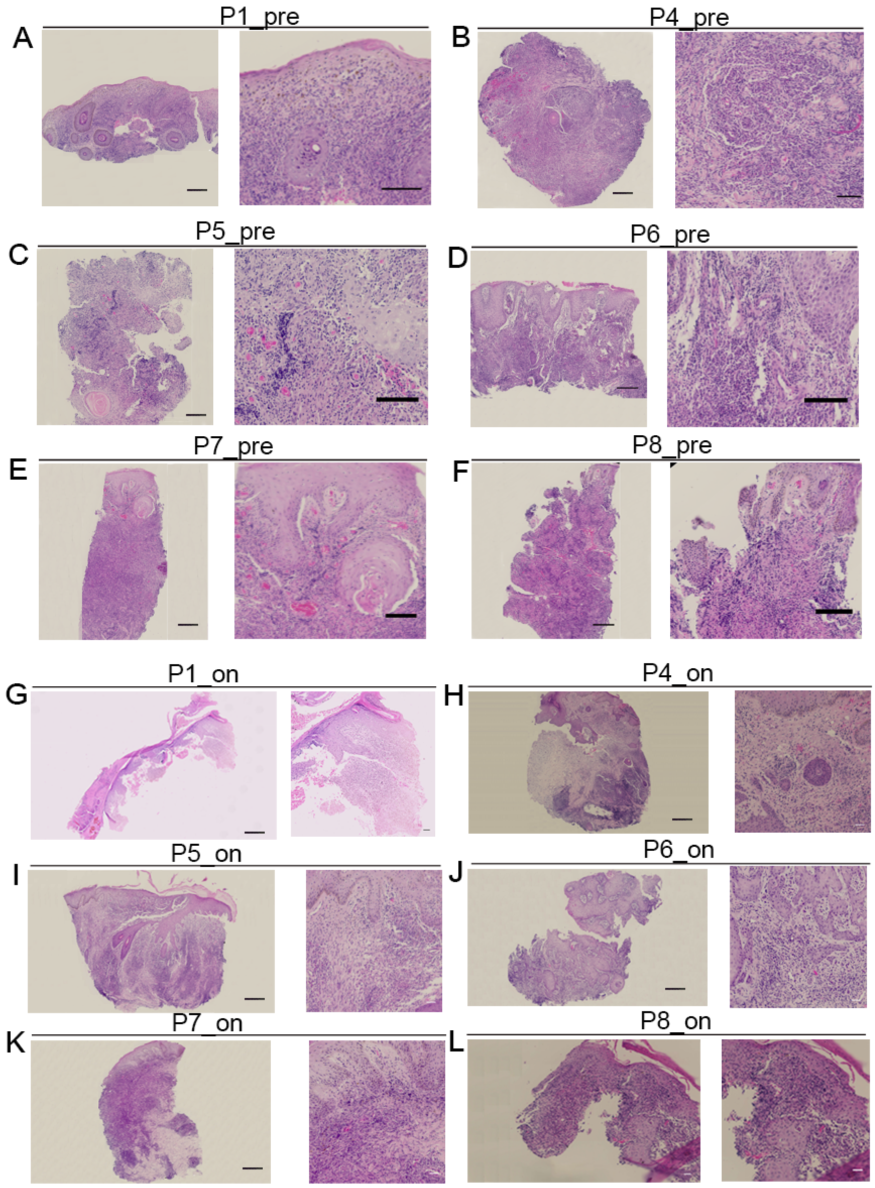


(A-L) Hematoxylin and eosin stains on whole 4um tissue sections and magnified areas from patients P1 (A), P4 (B), P5 (C), P6 (D), P7 (E) and P8 (F) at presentation and on treatment (G-L). Scale bar is 100u in whole sections and 20u in zoomed in areas.

**Supplemental Figure 4. RNAseq analysis of whole blood in CL infected patients**

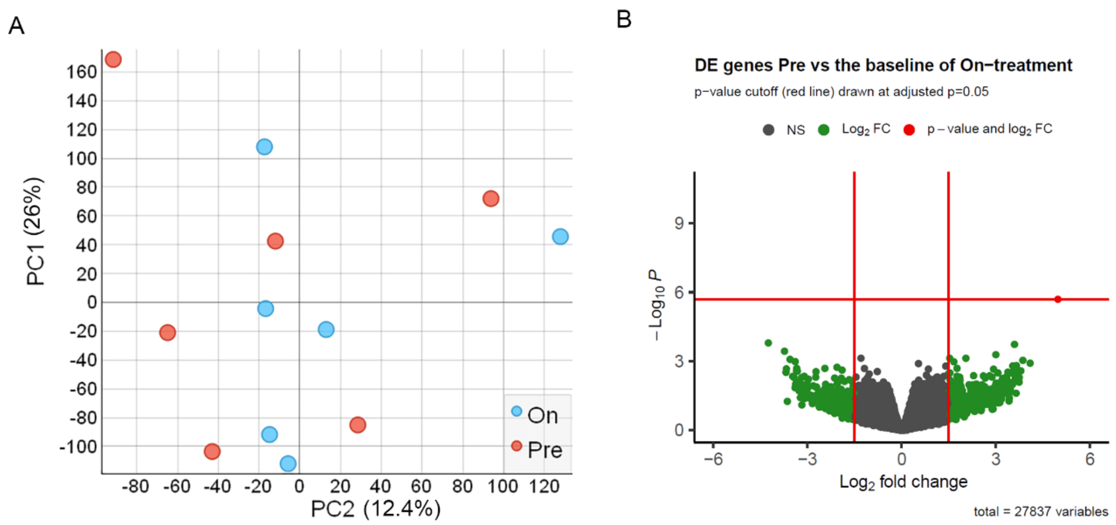


(A) Principal component analysis (PC1 vs PC2) on 27837 features from RNAseq dataset from whole blood of 6 CL patients (test cohort) at baseline (red dots) and after 2 weeks of treatment (red dots). (C) Volcano plot showing differentially expressed (DE) genes in CL patients at baseline relative to 2 weeks of treatment. Red lines are drawn at FDR = 5% and LogFC>1.5.

**Supplemental Figure 5. ROI strategy for DSP**

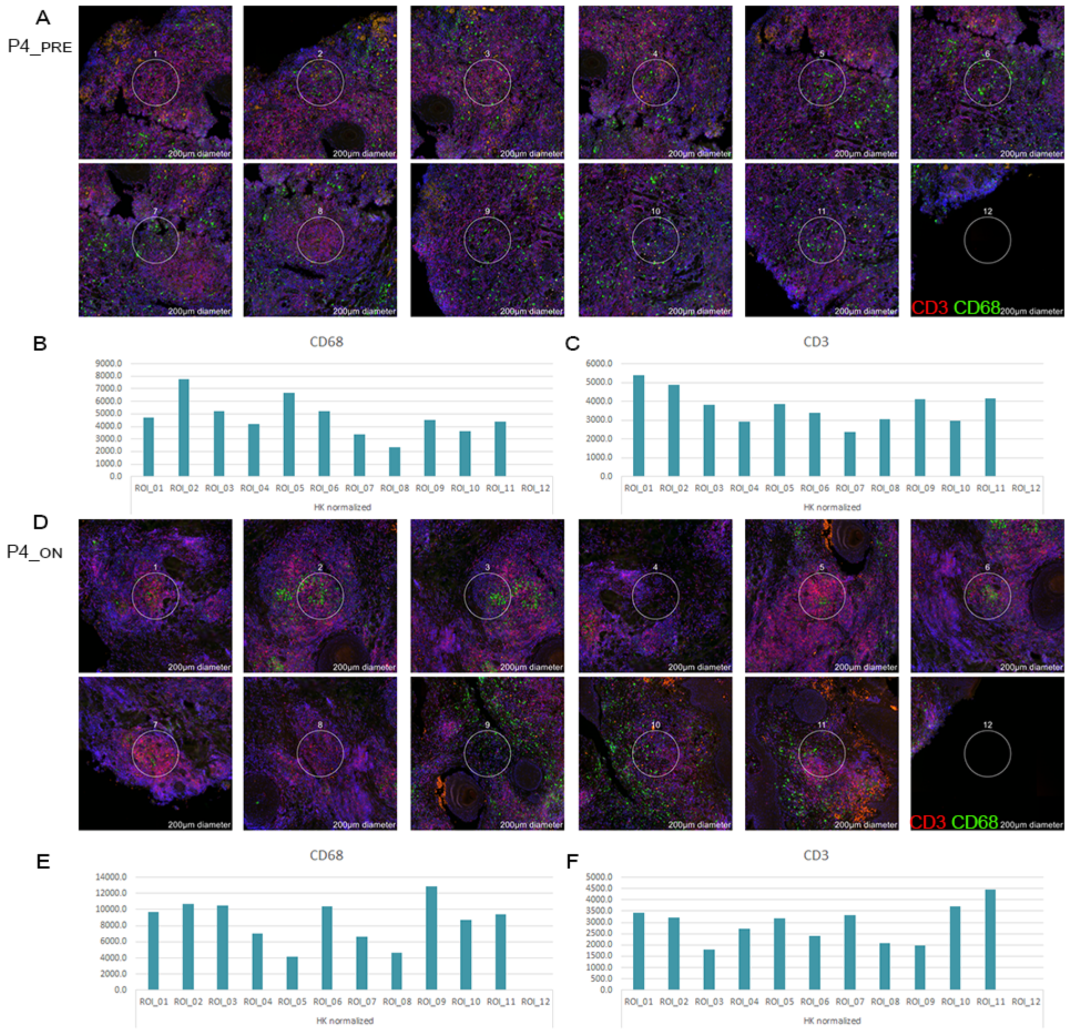


12 ROIs were generated pre-treatment and 12 ROIs were generated on-treatment for each patient. **(A)** Representative images from patient P4 of each ROI at presentation. ROIs were created on both $CD3^{+}$ and/or $CD68^{+}$ immune infiltrates, with a control ROI on glass (bottom right). **(B-C),** Bar graphs show housekeeping (HK) normalised CD68 (**B**) and CD3 (**C**) counts in each ROI. **(D)** Representative images from patient P4 of each ROI selected on-treatment. **(E-F)** Bar graphs show housekeeping (HK) normalised CD68 (**E**) and CD3 (**F**) counts in each ROI.

**Supplemental Figure 6. DSP: FDR passed discoveries and protein-protein interaction network**

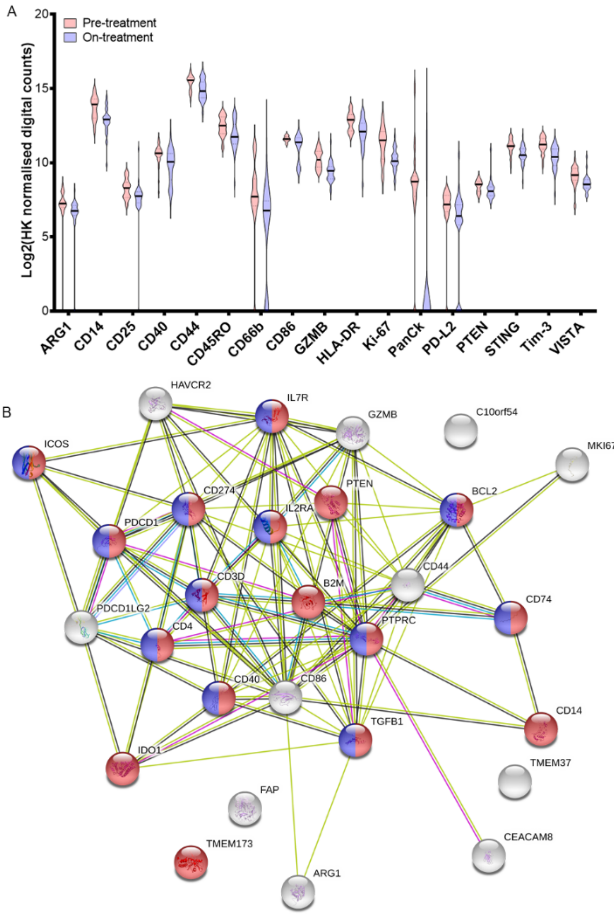


| #pathway ID | pathway description | observed gene count | false discovery rate | matching proteins in network |
|---|---|---|---|---|
| GO.0002684 | positive regulation of immune system process | 17 | 2.70E-14 | B2M,BCL2,CD14,CD274,CD3D,CD4,CD40,CD74,ICOS,IDO1,IL2RA,IL7R,PDCD1,PTEN,PTPRC,TGFB1,TMEM173 |
| GO.0050863 | regulation of T cell activation | 12 | 1.58E-13 | C10orf54,CD274,CD3D,CD4,CD74,ICOS,IDO1,IL2RA,IL7R,PDCD1,PTPRC,TGFB1 |

53 features were compared from 33ROIs each obtained from pre-treatment and on-treatment biopsies of 3 patients (P4, P6 and P7). Mann Whitney test was performed and corrected for FDR=5%. (A) Violin plot showing background corrected, HK normalised protein counts for discoveries comparing pre-treatment vs on-treatment ROIs (top 17 based on q value). (B) STRING db analysis of all discoveries (FDR = 5%)

**Supplemental Figure 7. Consort diagram of validation cohort patients**

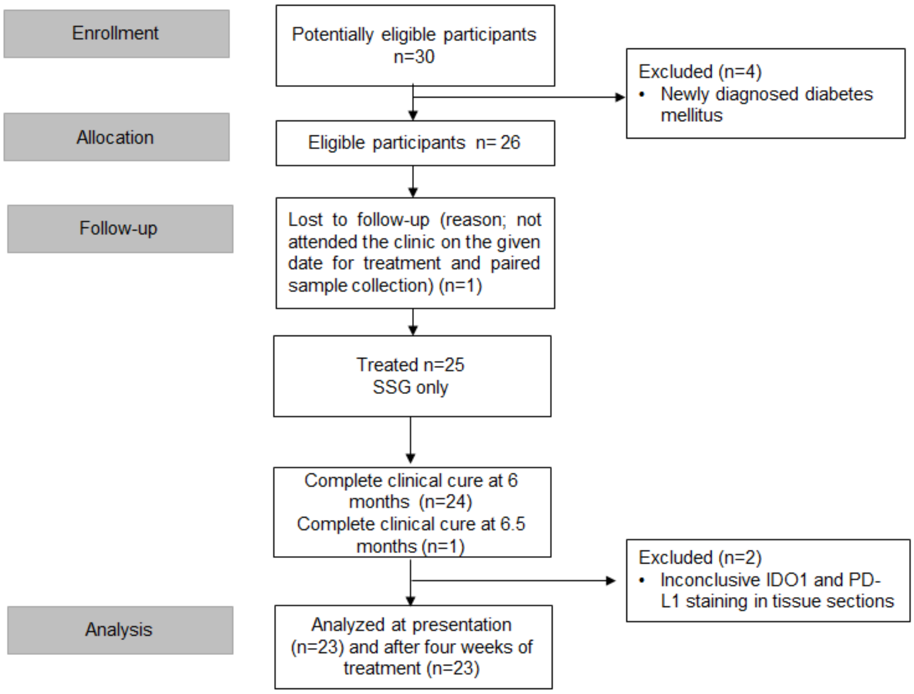


Flow chart shows number of patients who were screened, treated, followed up in early assessment, completed the study and included in the final analyses SSG: Sodium Stibogluconate; Cryo: Liquid nitrogen cryotherapy

**Supplemental Figure 8. Patient photographs and histology of individuals from validation cohort**

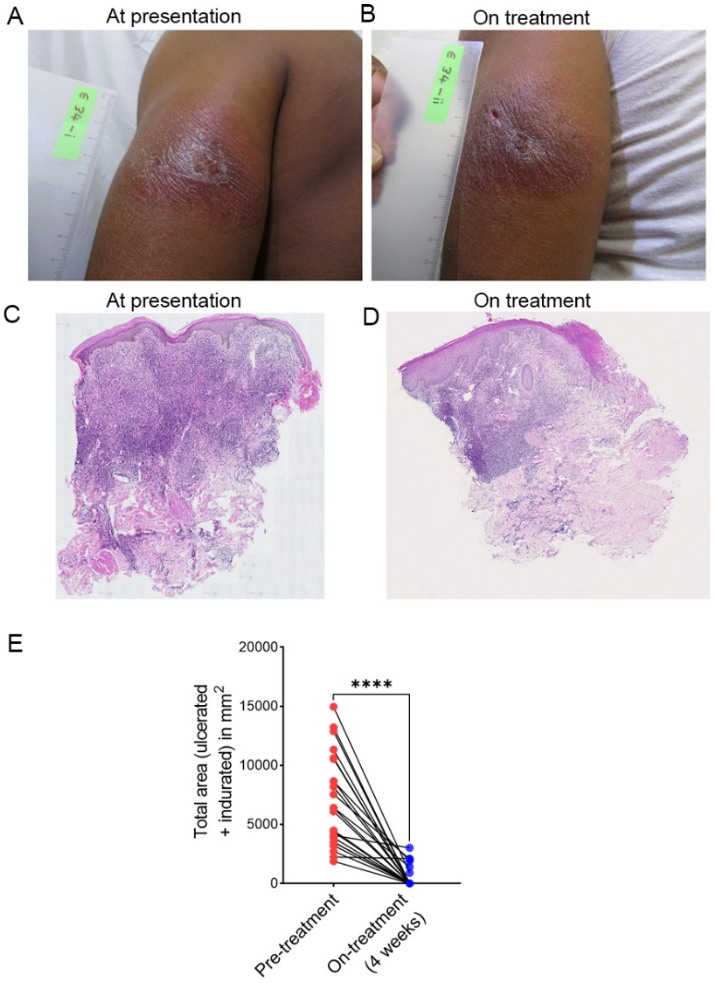


**A-C** Representative patient photograph at baseline (**A**) and showing improvement after 4 weeks (**B**). **(C-D),** H & E staining on tissue biopsy sections taken at presentation (**C**) and after 4 weeks of treatment (**D**). (**E**) Comparison of total area (ulcerated area and indurated area) per patient at presentation and after 4 weeks of treatment.

**Supplemental Figure 9. Gating strategy for low, medium and high PD-L1 expressing cells and Amastin expression**

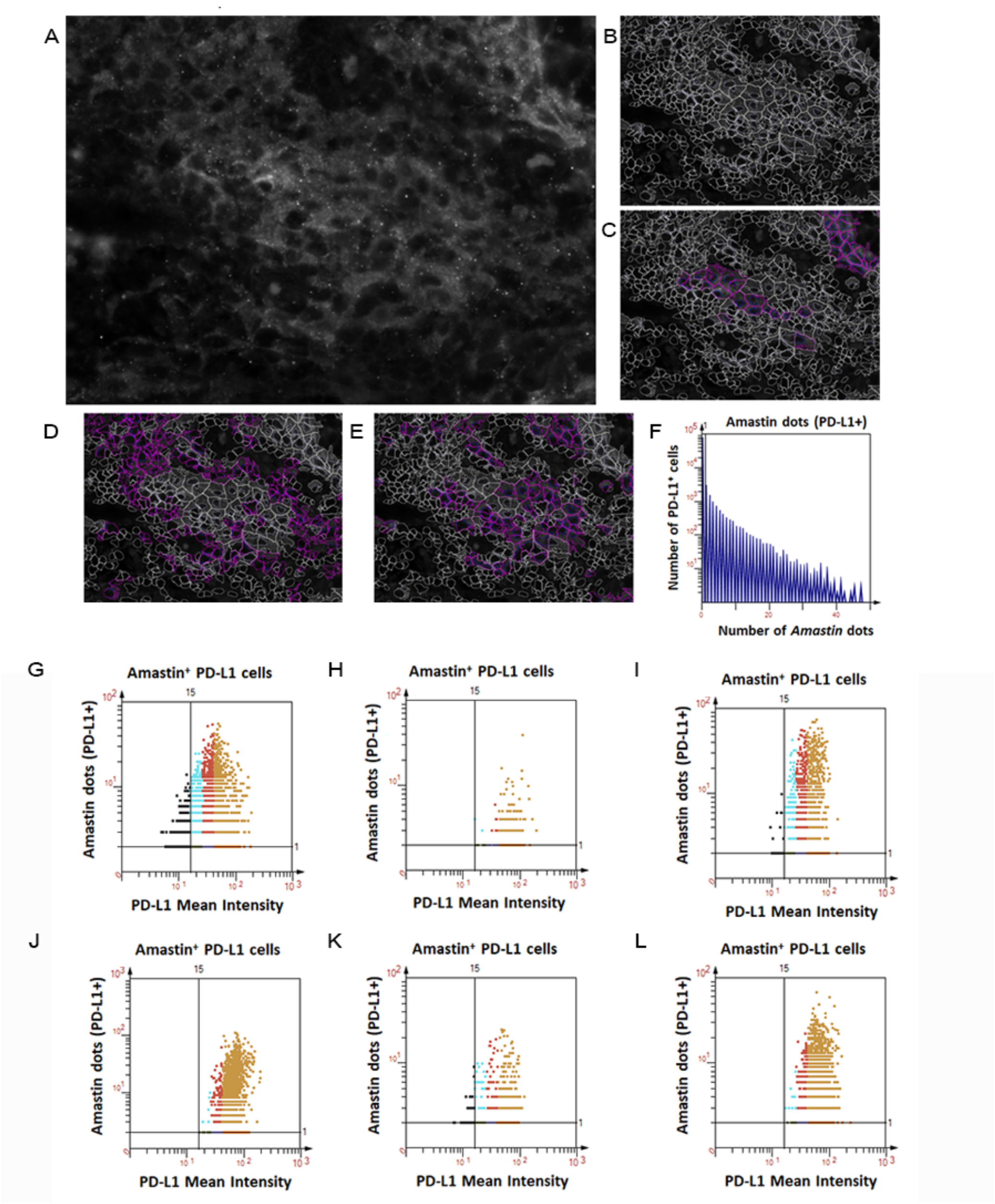


**A)** Grey image of magnified PD-L1 expression in skin tissue biopsy. (**B**) PD-L1$^{+}$ cells as detected using Strataquest image analysis software (**C-E**) Cells were binned based on mean fluorescence intensities of each cell into high (**C**), moderate (**D**) and low (**E**) expression, respectively. (**F**) Representative histogram showing distribution of *Amastin* dots in PD-L1$^{+}$ cells. Axis scale was adjusted to demonstrate the cut off (1 *Amastin* dot) used for PD-L1$^{+}$ cells containing *Amastin* dots. Some data points are off scale. (**G-L**) Scattergram from *Amastin*$^{+}$ cells at presentation from P9 (**G**), P13 (**H**), P16 (**I**), P17 (**J**), P23 (**K**) and P29 (**L**) respectively, showing number of *Amastin* dots in low (cyan), medium (red) and high (dark yellow) PD-L1 expressing cells.

**Supplemental Figure 10. Infected PD-L1$^+$ cells have greater fluorescence intensity than uninfected PD-L1$^+$ cells**

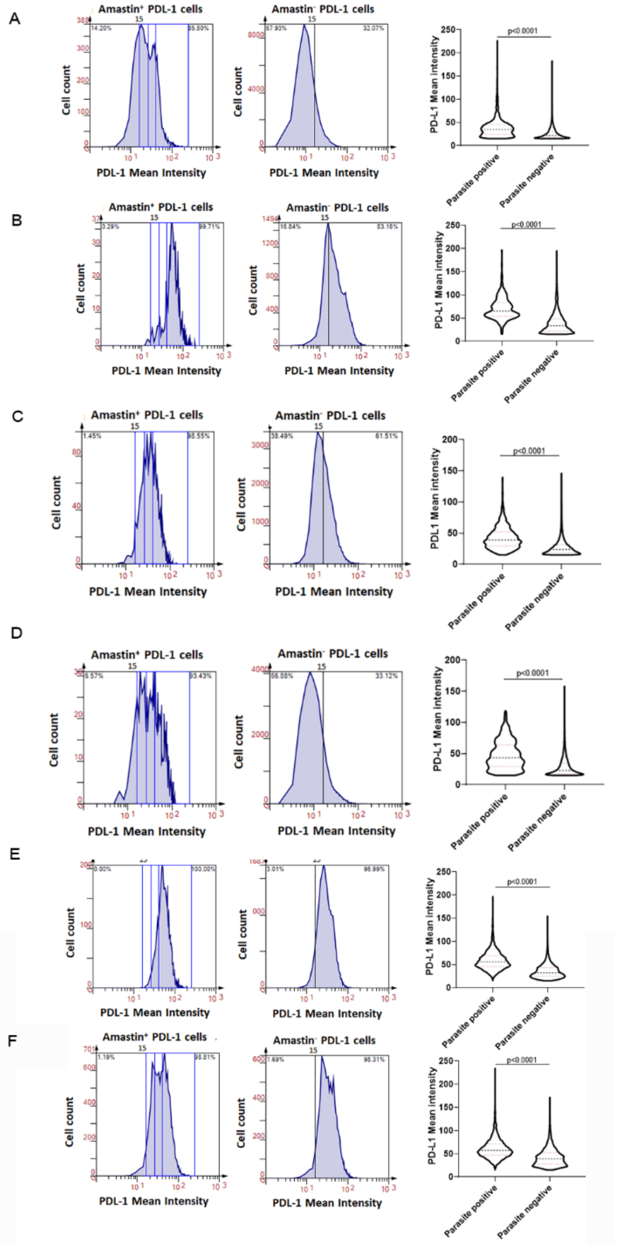


**(A-F)** Histograms and violin plots showing fluorescence intensity distributions of all infected and uninfected PDL1 cells of P9 (**A**), P13 (**B**), P16 (**C**), P17 (**D**), P23 (**E**) and P29 (**F**) at presentation respectively. Dotted lines show median, upper and lower quantile for each group. N>240 and >9000 were analysed for parasite positive and negative cells respectively from each patient.

**Supplemental Figure 11. PDL1 expression by hMDMs after in vitro infection with *L.donovani* zymodeme MON-37 or *L. major* Friedlin.**

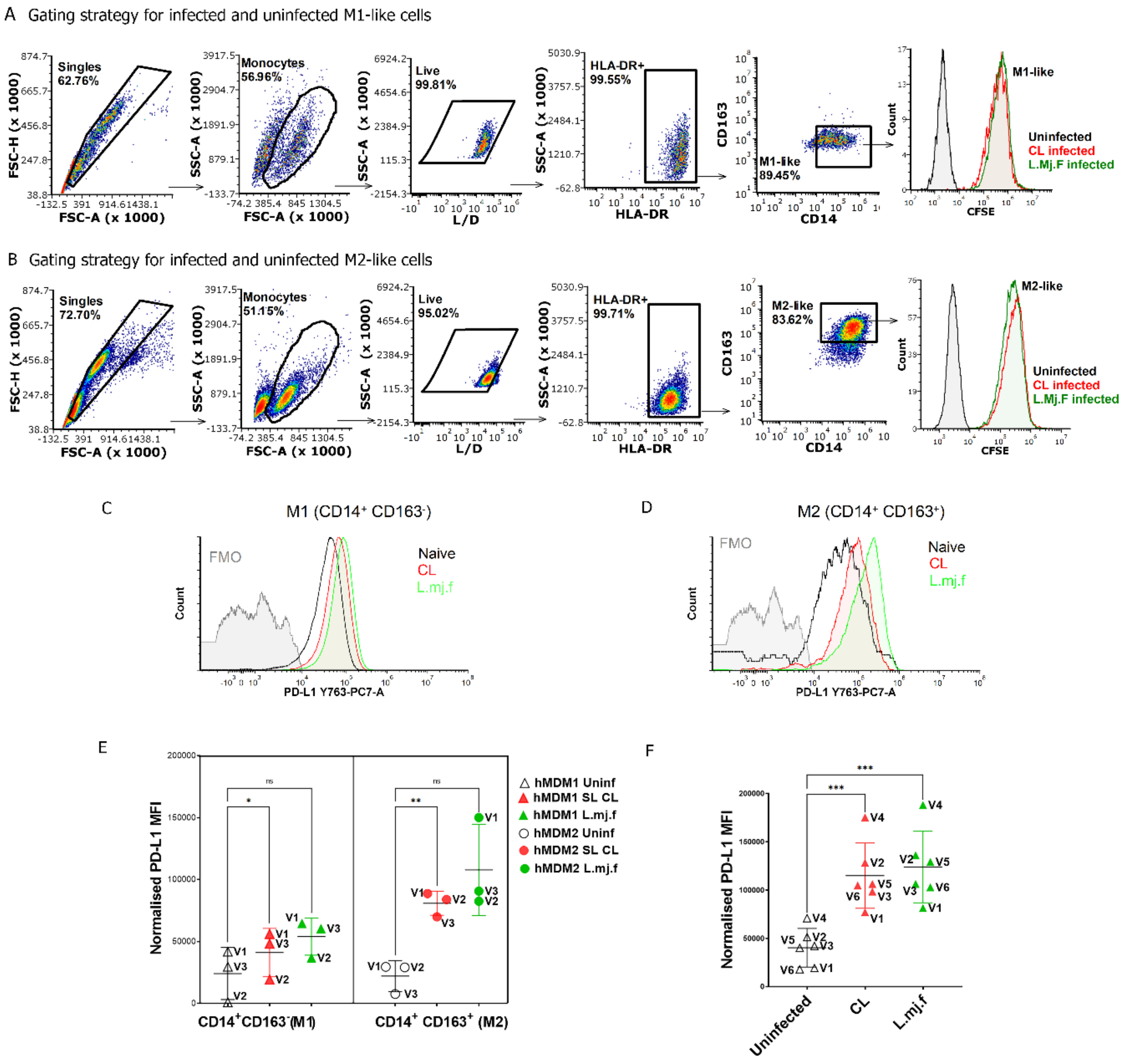


MACS-purified CD14$^{+}$ cells from PBMCs were differentiated using M-CSF (M1) or GM-CSF (M2) and either left uninfected or infected with *L.donovani* zymodeme MON-37 (CL) or *L.major* Friedlin (L.mj.f) stationary-phase promastigotes labelled with CFSE. (**A-B**) Gating strategy for M1 (HLADR$^{+}$ CD14$^{+}$ CD163$^{-}$) and M2 (HLADR$^{+}$ CD14$^{+}$ CD163$^{+}$) macrophages. (**C-D**) Representative histogram (from donor V1) showing PD-L1 expression (mean fluorescence intensity; MFI) on M1 (**C**) and M2 (**D**) uninfected macrophages (black histogram) and macrophages infected with CL (red) or L.mj.f (green) (**E**) Impact of infection on PD-L1 expression on uninfected M1 (left panel) and M2 macrophages (right panel) macrophages from 3 donors (V1-V3); white symbols, uninfected macrophages;

red symbols, CL-infected macrophages; green symbols, L.mj.F (**F**) PD-L1 expression in M1 an independent cohort of 6 volunteers (V1-6). Raw FCS files were analysed in FCS Express v7 and graphed in GraphPad Prism v9. Data are shown as mean ±SEM and were analysed using two-way RM ANOVA with Tukey's multiple comparisons test.

**Supplemental Figure 12. Intracellular parasitism in IDO1 expressing cells**

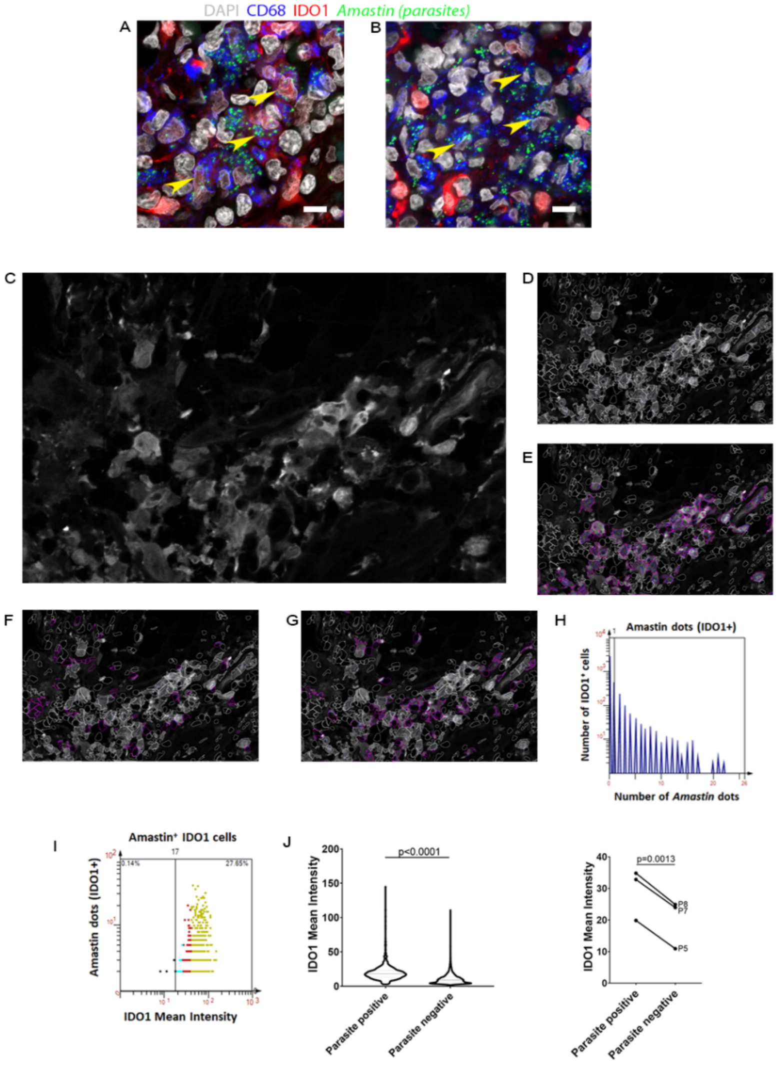


**(A-B)** Confocal images showing expression of IDO1 (red), CD68 (blue) proteins and *Amastin* RNA probe. Yellow arrowheads show infection in both IDO1$^{+}$CD68$^{+}$ (left panel) and IDO1$^{-}$ CD68$^{+}$ (right panel) cells. Scale bar, 10 pixels.Grey image of magnified heterogenous IDO1 expression in skin tissue biopsy. **(C-D)** IDO1$^{+}$ cells as detected using Strataquest image analysis software. **(E-G)** Cells were binned based on mean fluorescence intensities of each cell into high (**E**), moderate (**F**) and low (**G**) expression, respectively. (**H**) Representative histogram showing distribution of *Amastin* dots in IDO1$^{+}$ cells. *Amastin*$^{+}$ and *Amastin*$^{-}$ are defined by an *Amastin* count of 1, as shown by vertical bar. Some data points are off scale (**G-H**) **(I)** Representative scattergram (from patient P7 at presentation) colour coded to show low, medium and high IDO1 labelled cells in cyan, red and green with respect to *Amastin* dots. Each dot represents a single cell. (**J**) Representative violin plot (from patient P7) showing fluorescence intensity distributions in *Amastin*$^{+}$ and *Amastin*$^{-}$ cells. Dotted lines show median, upper and lower quantile for each group. N=791 and 3872 for parasite positive and negative cells, respectively. Significance score was generated using Students two tailed unpaired t-test. **(K)** Mean intensity of *Amastin*$^{+}$IDO1$^{+}$ cells when compared with *Amastin*$^{-}$ IDO1$^{+}$ cells (n=3 patients). Significance score was generated using Students two tailed paired t-test.

**Supplemental Figure 13. Univariate Cox Proportional Hazards model for PD-L1 and IDO-1 Kaplan-Meier plots**

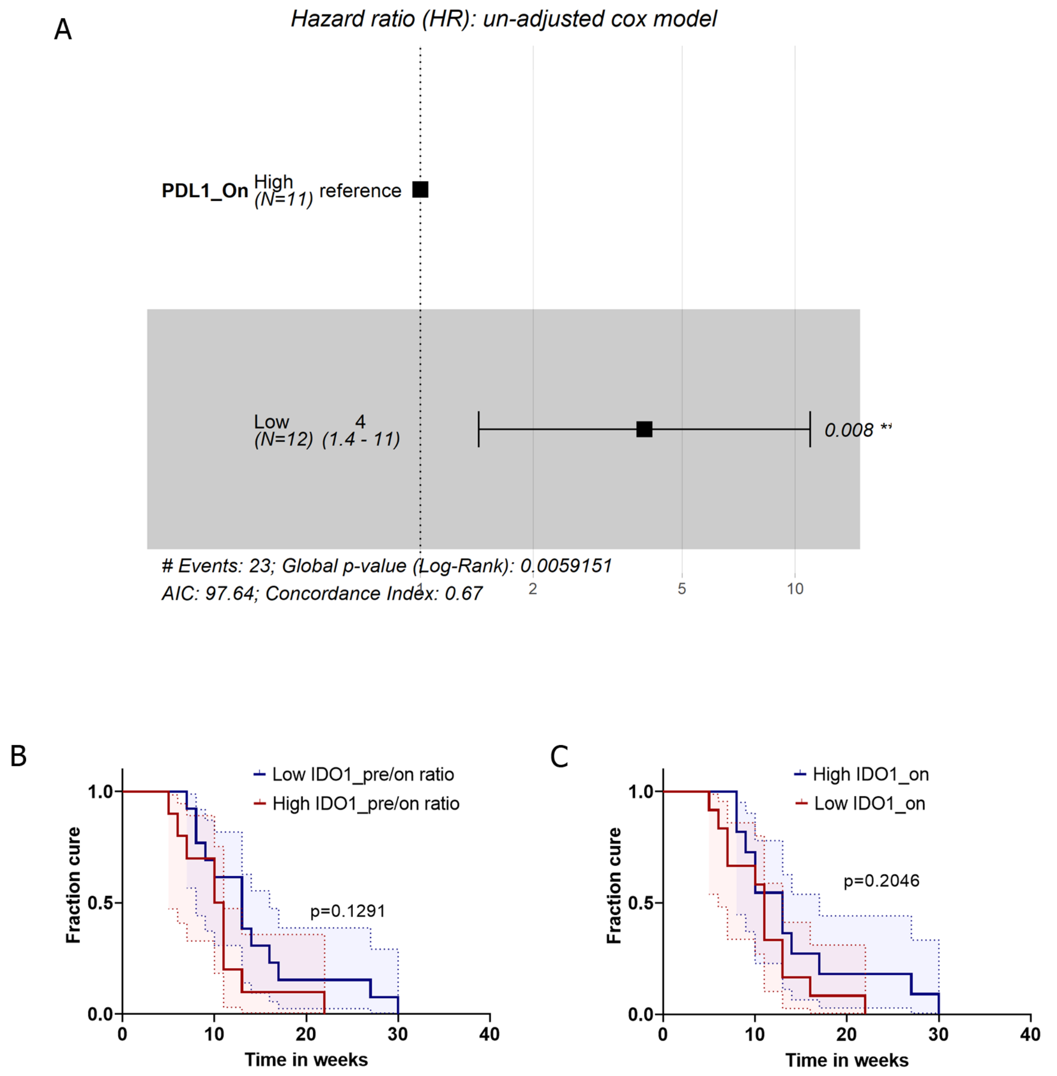


(**A**) Univariate Cox Proportional Hazard model plotted as a forest plot showing effect size of on-treatment expression of PD-L1 on cure rates. Hazards Ratio =3.96. p-values for PD-L1 and overall statistical significance is indicated and represent Wald statistic value. **B and C.** Patients (validation cohort; n=23) were stratified based on IDO1 expression at presentation and on therapy and cure rate plotted as a Kaplan-Meier curve. (B) Patients were stratified based on high (>geomean value; n=11) and low (< geomean value; n=12) pre-: on-treatment expression ratio. Kaplan-Meier curve based on pre:on ratio was plotted (C) Patients stratified based on-treatment expression of IDO1 (> geomean value; n= 11 vs < geomean value; n=12.
